# Identification of merozoite secreted repertoire and immuno-pharmacological inhibition of a novel host-parasite interaction to block malarial infection

**DOI:** 10.1101/2023.04.15.537002

**Authors:** Niharika Singh, Akshay Munjal, Geeta Kumari, Shikha Kaushik, Amandeep Kaur Kahlon, Sakshi Gupta, Ayushi Chaurasiya, Zill-e- Anam, Mukesh Kumar Maurya, Pallavi Srivastava, Jhalak Singhal, Manisha Marothia, Prerna Joshi, Ravi Jain, Devasahayam Arokia Balaya Rex, T. S. Keshav Prasad, Manoj Mundae, Pawan Malhotra, Anand Ranganathan, Shailja Singh

## Abstract

**Background:** During the intra-erythrocytic proliferation of *Plasmodium falciparum*, the host erythrocyte invasion is regarded as a complex and tightly regulated process comprising multiple receptor-ligand interactions, and numerous secretory molecules. Proteins secreted sequentially from apical organelles of merozoites serve as adhesins that play a crucial role in RBC invasion and can serve as vaccine and therapeutic targets.

**Methods:** Purified merozoites were triggered to discharge apical organelle contents by exposure to ionic conditions mimicking that of blood plasma. The secreted proteins were subjected to tandem mass spectrometry, and a well-characterized invasion ligand, RhopH3, was identified. A novel RhopH3 receptor, 14-3-3□ was unearthed using a Bacterial two-hybrid approach. This interaction was confirmed using multiple biophysical and biochemical approaches. We were successful in disrupting this interaction using a de novo peptide binder of 14-3-3□, and we subsequently assessed its effect on merozoite invasion.

**Results:** A total of 66 proteins were identified in the secretory fraction with apical organellar or merozoite membrane localization. The well-known adhesin, RhopH3 was also identified and its interaction with the host phosphopeptide-binding protein, 14-3-3□ was established. We also discovered a de novo peptide with the potency to disrupt this crucial interaction, thereby blocking merozoite invasion.

**Conclusion:** We, for the first time, report the secretory repertoire of plasmodium merozoite. Our study shows the importance of the erythrocyte protein, 14-3-3□ during the invasion process and paves the way for developing anti-malarial peptides or small molecules that inhibit the host-pathogen interaction, hence abrogating the invasion process.

## INTRODUCTION

Erythrocyte invasion by *Plasmodium falciparum* is indispensable for parasite survival and is crucial to malaria pathogenesis. The invasion of the erythrocyte begins with merozoite contact with the RBC membrane, followed by re-orientation to bring its apical end in close proximity to the RBC membrane^1, 2^. The apical end of merozoite contains apical organelles such as micronemes and rhoptries and harbor proteins that are released during the invasion.

Exposure of merozoites to low K^+^ ions (5 mM, which mimics the physiological ionic strength of human plasma) increases intracellular Ca^2+^ levels, triggering the secretion of micronemal proteins followed by rhoptry contents^3, 4^. Besides this, during the invasion, many MSPs are processed and shed in the extracellular milieu^5^. The repertoire of merozoite secretome is not currently available, and the direct encounter of these proteins with the host-immune system makes them a lucrative vaccination target. As a result, their identification could set up a platform for the development of subunit, chimeric vaccines, or other therapeutic interventions.

Using a proteomics approach, we attempted to characterize the proteins secreted by merozoites during host cell invasion. Surprisingly, we found a well-characterized Rhoptry resident adhesin, RhopH3 in the secreted fraction with high confidence. RhopH3 is essential for host cell invasion and nutrient uptake, and its C-terminus is conserved^6, 7^. RhopH3 has previously been shown to interact with cyclophilin B and Band 3 on the erythrocyte surface^8, 9^. Furthermore, the RhopH3 peptide or antibody inhibits merozoite invasion^9–11^. However, we anticipate that *Pf*RhopH3 interacts with a multiprotein complex on the erythrocyte, promoting merozoite binding and invasion. Thus, to identify a novel erythrocyte receptor of *Pf*RhopH3, we used the B2H approach to screen a human cDNA library against a C-terminal fragment of *Pf*RhopH3 as a bait, and we found human 14-3-3□ as an interacting partner of *Pf*RhopH3C. To further investigate the relevance of this binding, we discovered an 87 aa long peptide binder against host 14-3-3□ from a de-novo shuffled library that inhibits this interaction and prevents merozoite invasion.

Altogether, these findings point to a new host-pathogen interaction and the subsequent discovery of a small binder that inhibits this interaction, which could serve as a valuable therapeutic target for further research.

## METHODS

The human lung cDNA library was acquired from Stratagene (Cat: 982201) and di-codon Library was prepared as described previously^8, 12, 13^.

### Parasite culture and merozoite enrichment

*Pf*3D7 isolate was cultured and the merozoites were isolated according to standard protocols^3, 14^. The purity of merozoite preparation was >90%. Purified merozoites were resuspended in buffer mimicking plasma ionic conditions [Extracellular buffer (EC) buffer; 25mM HEPES (pH 7.2), 5.6mM glucose, 140mM NaCl, 5mM KCl, and 1mM CaCl_2_] and incubated at 37°C for 15 minutes, then separated from the secretory fraction. Secreted fractions and merozoite pellets were pooled from four independent experiments and subjected to LC-MS/MS for protein identification.

### In-solution digestion and LC-MS/MS analysis

20 µg of protein from the secretory and pellet fractions was reduced with 10mM DTT and alkylated with 20mM IAA. The proteins were digested with trypsin (1:20) at 37°C for 16 hours. The reaction was stopped by adding 0.1% formic acid, desalted with SCX, and the eluted peptides were lyophilized and stored. The peptides were analyzed using a Thermo Scientific Orbitrap Fusion Tribrid mass spectrometer connected to an Easy-nLC-1200 nano flow liquid chromatography system, as described previously^15^.

### Identification of peptides and proteins

Raw MS files were searched using Proteome Discoverer software suite version 2.2 (Thermo Fisher Scientific). *Pf3D7* protein reference database was retrieved from PlasmoDB, and MS/MS data were searched against the protein database as well as known mass spectrometer contaminates using the SEQUEST and Mascot algorithms. The search parameters that were employed were previously described^15^.

### Bacterial two-and three-hybrid assay

The experiment was done according to the standardized protocols of our lab^8, 12, 16, 17^. Briefly, plasmid pBTnn harboring the *Pf*RhopH3C coding sequence (bait) was screened against a Human lung cDNA library cloned into plasmid pTRGqq, and host 14-3-3□ was fished out as a potential binder. Similarly, the 14-3-3□ pTRGqq was used as bait and screened against the de novo KLRT pBTqq dicodon library, yielding a potential peptide binder NS1.

For bacterial three hybrid (B3H), the reporter strain R1 harboring both the bait (*Pf*RhopH3C-pBTnn) and prey (14-3-3□-pTRGqq) plasmids were co-transformed with the third plasmid, NS1-pMTSA, and plated on X-gal plates in the presence or absence of L-arabinose. The effect of pMTSA-encoded NS1 on the interaction of the other two proteins was assessed by reversion of colony color from blue to white. To quantify this interaction inhibition, a B3H arabinose gradient liquid β-galactosidase assay with increasing concentrations of L-arabinose (0 to 2mM) was performed.

### Cloning and purification of recombinant 14-3-3**□** (full length and TBD), *Pf*RhopH3-C, and NS1

*Pf*RhopH3 (C-terminal portion; 617-865 aa, 747 bp) and 14-3-3□ (full length; 246 aa, 738 bp) genes were codon-optimized, synthesized de novo, and subcloned into SnaB1 digested pMTSAra and pET-28a(+) vectors, respectively. 14-3-3□ TBD and NS1 constructs were PCR-amplified using specific primers and cloned into the pMTSAra vector at the SnaB1 restriction site. The expression of recombinant proteins was induced by L-Arabinose (for pMTSAra) and IPTG (for pET-28a(+)). Recombinant proteins were purified by Ni-NTA affinity chromatography. Rabbit polyclonal antisera were raised against RhopH3C.

### *In vitro* protein-protein interaction

All the experiments were performed following the standard protocols with minor modifications^8, 17–19^. For far western, 14-3-3□ full length or TBD were transferred onto PVDF membrane, incubated with r*Pf*RhopH3C, and detected using anti-RhopH3C. For ELISA, *Pf*RhopH3C or NS1 (200ng) were coated overnight on an ELISA microtiter plate, then blocked and incubated with either 14-3-3□ TBD or full-length protein. After washing, anti-14-3-3□ TBD (1:5000) and anti-14-3-3□ full length (1:10,000) were added, followed by incubation with the appropriate secondary antibody (1:5000) and O.D._450_ was measured. For ITC, a syringe was loaded with 14-3-3□ (78µM) which was then titrated with 5µM of RhopH3C maintained in the sample cell, and thermodynamic parameters were recorded. For MST, 14-3-3□ (10μM) was labeled with amine-reactive dye, NT-647 NHS (30µM), following the manufacturer`s protocol, and then titrated with different concentrations of *Pf*RhopH3C and NS1. Data analysis was performed with MO Affinity software^20^. For the pull-down assay, Ni-NTA bead-bound *Pf*RhopH3C (5μg) was incubated with 100μg of RBCs ghost lysate. Following washing, bound proteins were eluted with increasing concentrations of imidazole, and 14-3-3□ was detected by western blotting.

### *In-silico* data collection

RhopH3C chain C and 14-3-3□ structures were retrieved from the RCSB PDB Database; PDB IDs: 7MRW and 2C63, respectively^21, 22^. The structure of potential de novo binder NS1 was modeled using I-TASSER^23^. *In silico* protein-protein interactions between 14-3-3□/RhopH3C, and 14-3-3□/NS1 were analyzed using HADDOCK 2.4^24^. BV02 structural file (CID: 1087337) was retrieved from the NCBI PubChem database (https://pubchem.ncbi.nlm.nih.gov) and docked with 14-3-3□ using AutoDock4.2^25^.

### Immunofluorescence assay

RBCs were fixed with PFG and incubated with anti-14-3-3□ (1:500) before being incubated with Alexa-fluor 488 conjugated anti-mice antibody. The mounted smears were observed under a fluorescence microscope. Similarly, RBCs were incubated with NS1 peptide, washed, and then incubated with specific primary and secondary antibodies. After washing, cells were smeared on a glass slide and visualized. For co-binding studies, purified mature schizonts were incubated with r14-3-3□ (10μg), probed with specific antibodies, and images were acquired in Olympus FLUOVIEW FV3000 confocal microscope using Olympus cellSens dimensions imaging software.

### Erythrocyte invasion assay

To evaluate the effect of BV02 and NS1 on erythrocyte invasion, percoll purified *Pf*3D7 schizonts at 1% parasitemia and 2% hematocrit were incubated with varying concentrations of BV02, NS1, and anti-14-3-3□ under standard conditions. The rings were assessed (after 8 hours) by examining thin smears of each assay well and compared with untreated controls.

### Statistical analysis

Two-tailed Student’s t-test was used to calculate the statistical significance of data and error bars represent the mean ± SD of three independent experiments.

## RESULTS

### Identification and functional analysis of the secretory repertoire of merozoites

In an attempt to identify crucial merozoite proteins that are processed and secreted during RBC invasion, we subjected purified merozoites to EC buffer, and the proteins discharged from apical organelles were identified by tandem MS (Fig. 1a). The secretome fraction included 324 proteins, while the merozoite pellet had 1064 proteins which coincided with the data previously reported by our lab^26^. 289 proteins were shared between the secretome and pellet fractions. Since cytosolic enzymes and ribosomal proteins are prominent contaminants in global proteomic studies^27, 28^, they were excluded from further analysis. Furthermore, proteins derived from the nucleus and food vacuole were identified as being likely to be derived from damaged nuclei or contamination of food vacuole in the merozoite fraction and were excluded from further analysis.

**Fig. 1:**
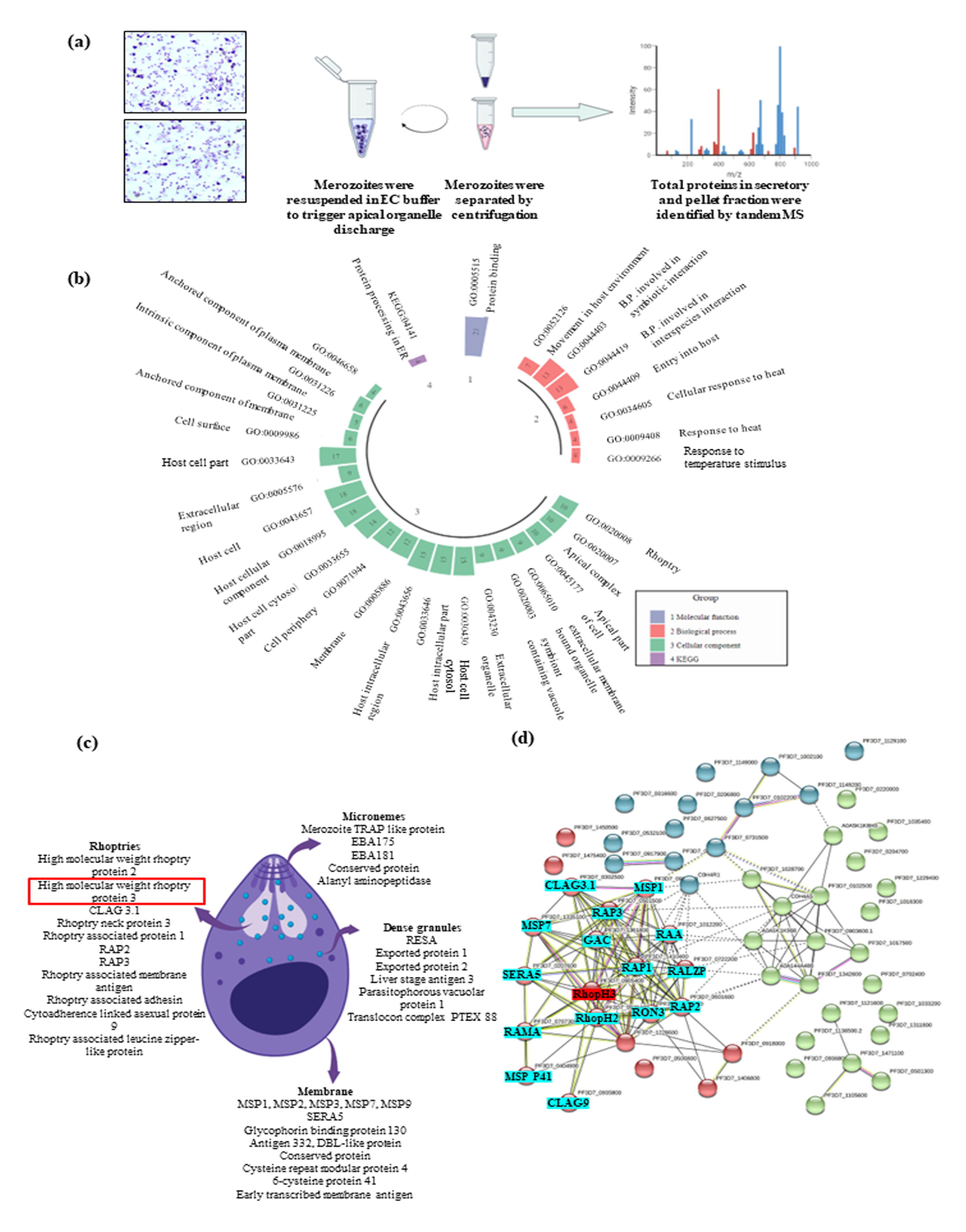
Identification and functional analysis of merozoite secretory repertoire. **a)** Proteomics approach used to enrich and identify proteins secreted by the merozoite during erythrocyte invasion. **b)** Gene ontology analysis of secreted proteins. **c)** Representation of key secretory proteins and their localization within the merozoite. **d)** Protein-protein interaction analysis of secretory proteins, highlighting the proteins implicated in merozoite invasion.

The remaining 66 identified proteins were grouped into eight different classes based on their subcellular localization in merozoite (as annotated by PlasmoDB database curators) (Table 1). To gain a functional understanding of the identified proteins, gene ontology analysis was performed using the software g:Profiler, as shown in Fig. 1b. Fig. 1c depicts a representation of key apical organelle and merozoite membrane proteins identified in our secretory fraction. The interactome of identified proteins was generated using the STRING database^29^, as shown in Fig. 1d. RAPs, MSP1,7 SERA5, CLAG3.1, RhopH2, CLAG9 and, RAMA were found in a major cluster of invasion-related proteins. RhopH3 was also identified in this interaction cluster, interacting with all of these proteins. We next attempted to identify a novel receptor on the erythrocyte surface that binds to RhopH3, resulting in the formation of a tight junction between the two cells that bridges them together.

**Table 1:** The proteins identified from proteomics analysis of merozoite secreted fraction. The predicted localization of each protein as predicted by PlasmoDB is indicated along with the number of peptides identified, its Molecular Weight (MW) and % coverage. The presence of Signal peptide (SP) or transmembrane domain (TMHMM) is also indicated for each protein.

| **PlasmoDB ID** | **Protein description** | **Predicted localization** | **Peptide Number** | **Molecular Weight (kDa)** | **Coverage (%)** | **Signal P or TMHMM** |
| --- | --- | --- | --- | --- | --- | --- |
| PF3D7_1311800 | alanyl aminopeptidase | Micronemes | 17 | 126 | 21 | TMHMM |
| PF3D7_0827900 | protein-disulphide isomerase | Micronemes | 10 | 55.5 | 34 | SP+<br>TMHMM |
| PF3D7_1028700 | merozoite TRAP like protein | Micronemes | 1 | 58 | 4 | SP+<br>TMHMM |
| PF3D7_1136500 | casein kinase 1 | Micronemes | 1 | 37.6 | 4 | - |
| PF3D7_0731500 | EBA175 | Micronemes | 1 | 174.5 | 1 | SP+<br>TMHMM |
| PF3D7_0903600 | conserved protein, | Micronemes | 1 | 90.6 | 2 | - |
| PF3D7_0102500 | EBA-181 | Micronemes | 20 | 181 | 20 | SP+<br>TMHMM |
| PF3D7_0930300 | MSP1 | Membrane | 19 | 195.6 | 15 | SP+<br>TMHMM |
| PF3D7_0206800 | MSP2 | Membrane | 3 | 27.9 | 24 | - |
| PF3D7_1016300 | Glycophorin binding protein 130 | Membrane | 2 | 95.8 | 32 | TMHMM |
| PF3D7_1335100 | MSP7 | Membrane | 2 | 41.3 | 8 | SP |
| PF3D7_0207600 | serine repeat antigen 5 | Membrane and extracellular vesicles | 2 | 111.7 | 2 | SP |
| PF3D7_0204700 | Hexose transporter | Membrane | 1 | 56.4 | 3 | TMHMM |
| PF3D7_1228600 | MSP9 | Membrane | 1 | 86.6 | 2 | SP |
| PF3D7_1035400 | MSP3 | Membrane | 1 | 40.1 | 5 | SP |
| PF3D7_1475400 | cysteine repeat modular protein 4 | Membrane | 1 | 700.5 | ND | SP+<br>TMHMM |
| PF3D7_1450500 | conserved protein | Membrane | 1 | 199.6 | ND | TMHMM |
| PF3D7_0532100 | Early transcribed membrane protein 5 | Membrane | 1 | 19 | 7 | SP+<br>TMHMM |
| PF3D7_0404900 | 6-cysteine protein 41 | Membrane | 1 | 43.1 | 3 | SP |
| PF3D7_1342600 | myosin A | Membrane | 8 | 92.2 | 14 | - |
| PF3D7_0316600 | formate nitrite transporter | Membrane | 1 | 34.4 | 3 | TMHMM |
| PF3D7_0929400 | high molecular weight<br>rhoptry protein 2 | Rhoptries | 25 | 162.6 | 22 | SP |
| PF3D7_0905400 | high molecular weight<br>rhoptry protein 3 | Rhoptries | 17 | 104.8 | 25 | SP |
| PF3D7_0302500 | CLAG3.1 | Rhoptries | 13 | 167.1 | 12 | SP |
| PF3D7_1252100 | rhoptry neck protein 3 | Rhoptries | 7 | 263 | 4 | SP+<br>TMHMM |
| PF3D7_1410400 | rhoptry associated<br>protein 1 | Rhoptries | 19 | 90 | 32 | SP |
| PF3D7_0501600 | rhoptry associated<br>protein 2 | Rhoptries | 6 | 46.7 | 19 | SP |
| PF3D7_0501500 | rhoptry associated<br>protein 3 | Rhoptries | 4 | 47 | 13 | SP+<br>TMHMM |
| PF3D7_0707300 | rhoptry associated<br>membrane antigen | Rhoptries | 4 | 103.6 | 8 | SP |
| PF3D7_1012200 | rhoptry associated<br>adhesin | Rhoptries | 2 | 31.8 | 9 | SP |
| PF3D7_0935800 | cytoadherence linked<br>asexual protein 9 | Rhoptries | 1 | 160.3 | 1 | SP+<br>TMHMM |
| PF3D7_0722200 | rhoptry associated<br>leucine-zipper like<br>protein | Rhoptries | 1 | 87.8 | 1 | SP+<br>TMHMM |
| PF3D7_0102200 | RESA | Dense<br>granules | 15 | 126.4 | 17 | - |
| PF3D7_1121600 | exported protein 1 | Dense<br>granules | 3 | 17.3 | 12 | SP+<br>TMHMM |
| PF3D7_1471100 | exported protein 2 | Dense<br>granules | 2 | 33.4 | 9 | SP |
| PF3D7_0220000 | liver stage antigen 3 | Dense<br>granules | 1 | 175.6 | 1 | TMHMM |
| PF3D7_1129100 | parasitophorous<br>vacuolar protein 1 | Dense<br>granules | 2 | 51.9 | 4 | SP |
| PF3D7_1105600 | translocon component<br>PTEX88 | Dense<br>granules | 1 | 90.7 | 1 | SP |
| PF3D7_1017500 | myosin essential light<br>chain ELC | Cell surface | 1 | 15.7 | 7 | - |
| PF3D7_1361800 | glideosome associated<br>connector | Cell surface | 12 | 290.8 | 6 | - |
| PF3D7_0627500 | protein DJ-1 | Cell surface | 3 | 20.3 | 22 | - |
| PF3D7_0918000 | glideosome associated protein 50 | IMPC | 1 | 44.6 | 2 | TMHMM |
| PF3D7_1406800 | glideosome associated protein with multiple membrane spans3 | IMPC | 1 | 32.9 | 3 | TMHMM |
| PF3D7_0917900 | Hsp70 | Cell surface, Extracellular vesicles | 18 | 72.3 | 38 | SP |
| PF3D7_1149200 | Ring infected erythrocyte surface antigen | Maurer's cleft | 5 | 126.8 | 3 | - |
| PF3D7_0501300 | skeleton binding protein 1 | Maurer's cleft | 1 | 36.3 | 7 | TMHMM |
| PF3D7_1033200 | early transcribed membrane protein 10.2 | Maurer's cleft | 1 | 38.9 | 5 | SP+ TMHMM |
| PF3D7_1129100 | parasitophorous vacuolar protein 1 | Dense granule, Maurer's cleft | 2 | 51.9 | 4 | SP |
| PF3D7_1002100 | EMP-1 trafficking protein | Maurer's cleft | 1 | 69.8 | 3 | TMHMM |
| PF3D7_0936800 | PHISTc | Maurer's cleft | 1 | 45.5 | 3 | TMHMM |
| PF3D7_1149000 | antigen 332, DBL-like protein | Maurer's cleft | 1 | 688.9 | 1 | - |
| PF3D7_1222300 | Endoplasmin | Extracellular vesicle | 20 | 95 | 31 | SP |
| PF3D7_0708400 | Hsp90 | Extracellular vesicle | 16 | 86.1 | 22 | - |
| PF3D7_0922200 | S-AM synthetase | Extracellular vesicle | 14 | 44.8 | 54 | - |
| PF3D7_1105000 | Histone H4 | Extracellular vesicle | 6 | 11.4 | 51 | - |
| PF3D7_0831700 | Hsp 70 | Extracellular vesicle | 7 | 75 | 11 | SP+ TMHMM |
| PF3D7_0202500 | Early transcribed membrane protein 2 | Extracellular vesicle | 1 | 11.5 | 11 | TMHMM |
| PF3D7_1343000 | Phosphatidyl ethanolamine transferase | Extracellular vesicle | 16 | 31 | 82 | - |
| PF3D7_0702400 | Small exported membrane protein 1 | Extracellular vesicle | 1 | 14.2 | 13 | TMHMM |
| PF3D7_0808600 | *PfEMP1* | Knobs | 1 | 256.4 | ND | - |

### Identification and functional analysis of merozoite total proteome

A total of 1090 merozoite proteins were identified in the pellet fraction, with a high confidence. These proteins were subjected to Gene Ontology analysis and were classified based upon their cellular component, molecular function, and biological processes in which they are involved. Housekeeping proteins such as Ribosomal proteins, nuclear proteins, metabolic enzymes, cytoskeletal proteins were abundantly detected in the total proteome. Besides this many membranes and apical organellar proteins that mediates host cell attachment and invasion were markedly detected. This includes the Merozoite surface proteins (MSP 5, 6, 7, 8, MTRAP, MSA 180) the rhoptry resident proteins (RAP1, RAP2, RAP3, RON2, RON4, RON5, RON6, RON11, RON14, RAMA, RALZP, RAMA, Rhop2, Rhop3, CLAG 3.1, CLAG3.2, CLAG9) and the micronemal proteins (EBA175, EBA181, AMA-1).

Also, 8.5% of the total proteins identified were annotated to be conserved with no known function (or hypothetical proteins). A comparison of our merozoite proteome with other published proteomes ^30^, revealed an overlap of greater than 46% (Fig. 2a). 585 proteins were unique to our proteins, whereas 335 were exclusive to Florens et al. These differences can be attributed changes in experimental and analytical techniques utilized in these studies. A gene ontology analysis of merozoite proteome was also carried out and the top enriched term IDs for the categories; Molecular function, Cellular components and Biological process is presented in Fig. 2 b, c, and d.

**Fig. 2:**
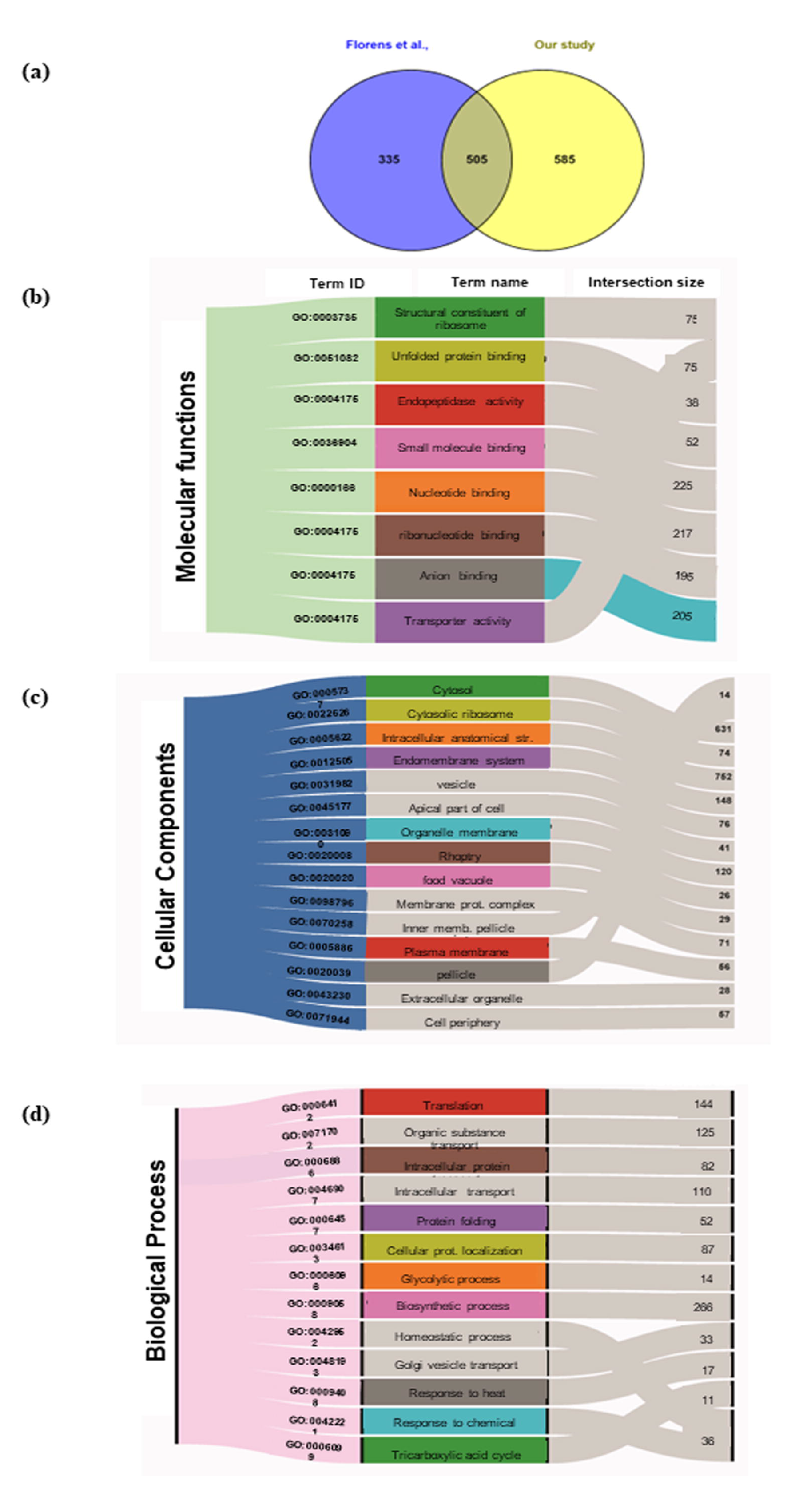
Gene ontology analysis of merozoite total proteome. a) Venn-diagram depicting the exclusive and overlapping proteins, between previously published merozoite proteomes and our study. b) Gene ontology enrichment analysis of protein identified in the merozoite pellet. The top enriched terms for the categories of molecular functions, cellular components and biological process are represented in (b-d). The intersection size of each term name is also given.

### Host 14-3-3**□** as a novel *Pf*RhopH3C interaction partner

Screening of the human cDNA library against the RhopH3 C-terminal region using the B2H approach identified human 14-3-3□ (gene ID: YWHAH) as the most likely host interaction partner (Fig. 3a). BLAST analysis indicated that this protein was not full-length 14-3-3□, but rather only the target binding domain (TBD), which is required for 14-3-3□ to bind to its targets^31^. Key interaction residues of *Pf*RhopH3C and 14-3-3□ are represented in Fig. 3b (i). A schematic representation of both proteins is depicted in Fig. 3b (ii). IFA indicated that 14-3-3□ localizes towards the cell periphery (Fig. 3c (i)). Western blotting corroborated these findings, detecting 14-3-3□ (∼28 kDa) in both the membrane and cytosolic fractions of RBCs (Fig. 3c (ii)). RhopH3 was found at the apical end of the merozoite, and native RhopH3 (∼110 kDa) was found in the schizonts lysate (Fig. 3d (i) and (ii)). To confirm if 14-3-3□ serves as a ligand for RhopH3, we assessed its binding to free merozoites. 14-3- 3□ was found to co-localize with *Pf*RhopH3, indicating a direct interaction of the two proteins (Fig. 3e (i)). The interaction was further validated by the pull-down assay in which native 14-3-3□ was pulled out from RBC lysate using rRhopH3C as bait (Fig. 3e (ii)). These findings suggested that 14-3-3□ could be a significant interaction partner of *Pf*RhopH3C during the invasion process.

**Fig. 3:**
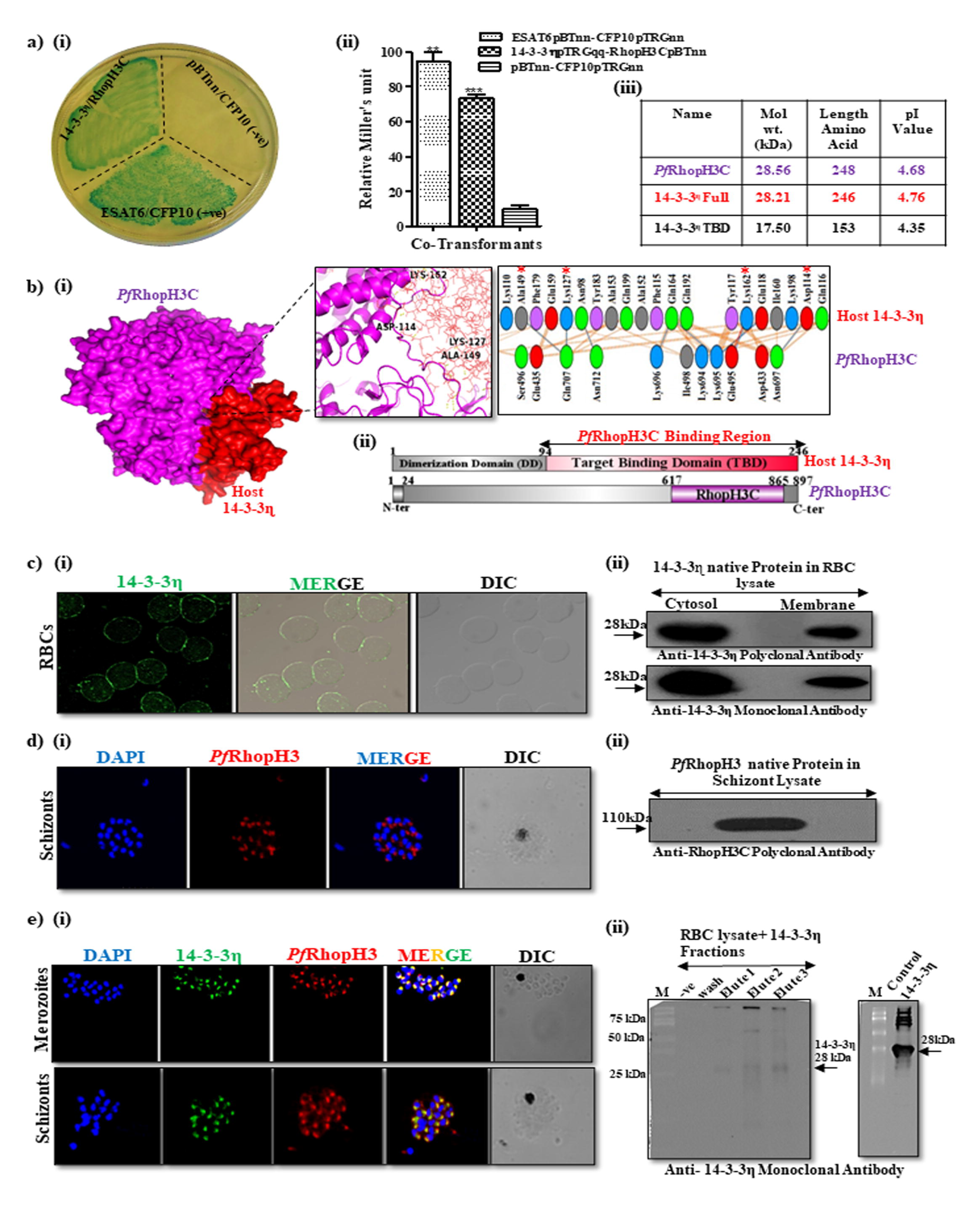
Human 14-3-3□ as a novel *Pf*RhopH3 interaction partner. **a) (i)** Screening of the human cDNA library against the RhopH3 C-terminal region using the B2H approach identified human 14-3-3□ as the interaction partner. Blue colonies indicated protein-protein interaction, and white colonies indicated no interaction. Test/ Negative/ Positive: RhopH3CpBTnn-14-3-3□pTRGqq/CFP10pTRGnn-empty pBTnn/CFP10pTRGnn-ESAT6pBTnn. **(ii)** β-galactosidase assay demonstrating enzymatic activity (in Miller’s Unit, MU) of the co-transformant pairs. **(iii)** Molecular parameters of proteins. **b) (i)** Surface view of the *Pf*Rhoph3C (purple)/14-3-3□ (Red) docked complex. **(ii**) Schematic of the overall architecture of 14-3-3□ and *Pf*RhopH3; 14-3-3□: 1-93 aa Dimerization Domain (DD), 94- 246 aa Target Binding Domain (TBD); *Pf*RhopH3: 1-24 aa signal sequence, 617-865 aa *Pf*RhopH3-C terminal fragment chosen for the present study. **c) (i)** IFA showing localization of 14-3-3□ on RBCs. **(ii**) Western blot analysis of native 14-3-3□ in fractionated RBC lysate. **d) (i)** IFA showing *Pf*RhopH3 at the apical end in late schizonts. **(ii)** Western blot analysis of native *Pf*RhopH3 in the schizont lysate showing the desired band at ∼110 kDa. **e) (i)** IFA showing binding of r14-3-3□ to mature schizonts and free merozoites. **(ii)** Host 14-3-3□ pull-down assay from erythrocyte lysate using resin-bound r*Pf*RhopH3C as bait. Wash: 0mM imidazole; elutes 1-3: elution fractions containing 10, 50 and 75mM imidazole; -ve: negative control. As a positive control, 14-3-3□ from RBC lysate was probed.

### Characterization of the 14-3-3**□** and *Pf*RhopH3C interaction

We confirmed the direct interaction of both proteins using multiple biochemical and biophysical approaches. ELISA indicated a concentration-dependent increase in OD_450_ with an increase in the concentration of overlaid 14-3-3□ (full length and TBD) confirming the interaction between RhopH3 and 14-3-3□ (Fig. 4a). ITC was performed to quantify the interaction affinities. The binding isotherms upon titration of RhopH3C with 14-3-3□ are shown in Fig. 4b and a binding affinity constant (K_a_) was determined to be 1µM. Furthermore, far-western analysis validated the interaction in which distinct bands corresponding to 14-3-3□ full length (28 kDa) and TBD (17 kDa) were observed, depicting a substantial interaction between the two proteins (Fig. 4c). The strength of this interaction was further assessed by MST, in which the dose-response curve from titration of labeled 14-3-3□ with different concentrations of RhopH3C depicted a dissociation constant (K_d_) of 66 nM (Fig. 4d), suggestive of a strong interaction between the two proteins.

**Fig. 4:**
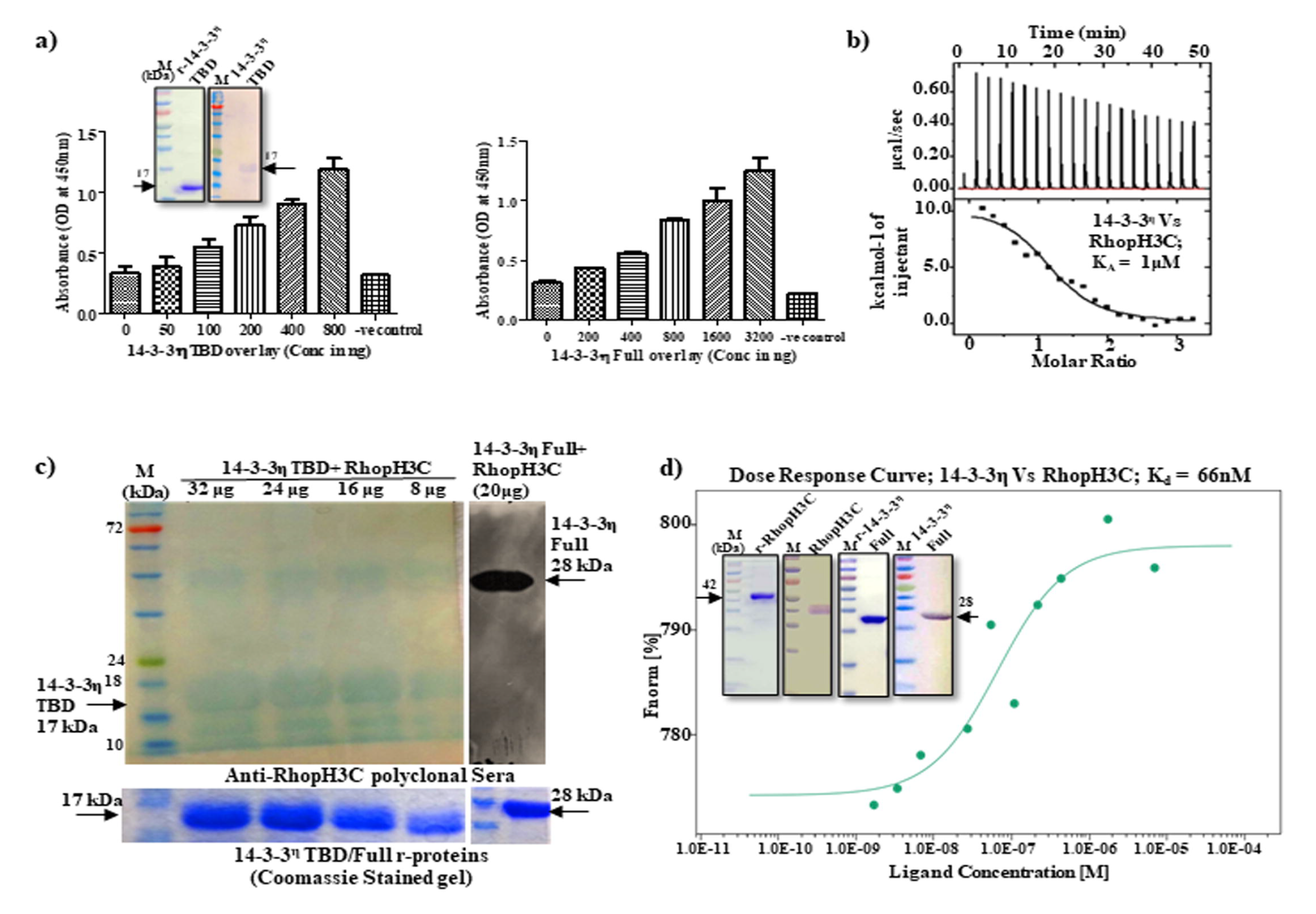
Characterization of 14-3-3□-*Pf*RhopH3C interaction. **a)** ELISA based interaction analysis of 14-3-3□ (full length/TBD) and *Pf*RhopH3C. **b)** Binding isotherms upon titration of 14-3-3□ (78 µM) with RhopH3C (5 µM). **c)** Far-western blot analysis demonstrating the desired bands of 14-3-3□ full length (28 kDa) and TBD (17 kDa). **d)** Dose-response curve resulting from MST between labeled 14-3-3□ and varying concentrations of RhopH3C.

### NS1 as a crucial disruptor for 14-3-3**□**-*Pf*RhopH3C interaction

After establishing the interaction of *Pf*RhopH3 with the host 14-3-3□, we attempted to disrupt this interaction using a novel short peptide. Using the B2H approach, 87 aa long peptide; NS1 was identified from an in-house generated di-codon library. The binding of 14-3-3□ with NS1 was assessed by co-transforming 14-3-3□-pTRGqq and NS1-pBTnn in R1 reporter cells and plating them on an X-gal plate (containing arabinose). The presence of blue colonies confirmed the interaction between 14-3-3□ and NS1 (Fig. 5a (i) and (ii)). 3D structure of NS1 is shown in Fig. 5a (iii). Recombinant NS1 was overexpressed in *E. coli* and purified to homogeneity (Fig. 5a (iv)). To further confirm the interaction between 14-3-3□ and NS1, ELISA was performed, in which a concentration-dependent binding between NS1 and 14-3-3□ was observed (Fig. 5b). K_d_ was determined to be 36 nM with MST, suggestive of a high affinity of the peptide for the protein (Fig. 5c). Erythrocyte binding assay revealed the potential of rNS-1 to bind 14-3-3□ on the RBC membrane (Fig. 5d). The B3H assay confirmed the disruption of 14-3-3□/*Pf*RhopH3C interaction by NS1 in the presence of L-arabinose, resulting in the conversion of blue colonies to white (Fig. 5e (i)). Whereas, in the absence of arabinose, the colonies remained blue (Fig. 5e (ii)). The arabinose gradient liquid β-galactosidase assay and inhibitory ELISA using NS1 established the concentration-dependent disruption of 14-3-3□/*Pf*RhopH3C interaction by NS1 (Fig. 5f and g).

**Fig. 5:**
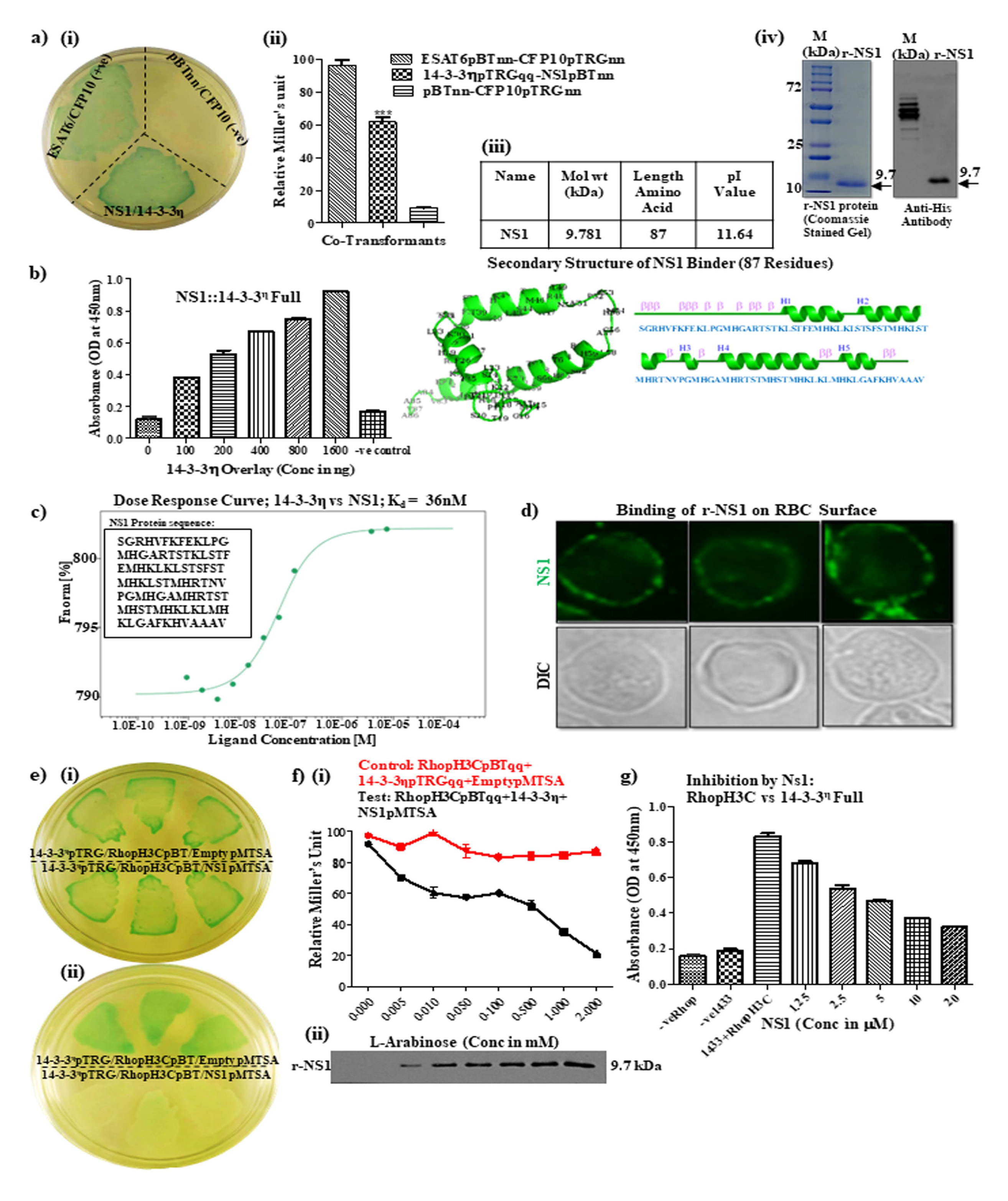
Identification of de novo binder against 14-3-3□ and disruption of 14-3-3□- *Pf*RhopH3C interaction. **a) (i)** Using the B2H approach, 87 aa long peptide NS1 was identified as the potential binder of 14-3-3□ from an in-house generated di-codon library. Blue colonies indicated the interaction between NS1-pBTnn and 14-3-3□-pTRGqq. **(ii)** β-galactosidase assay demonstrating enzymatic activity (in MU) of the co-transformant pairs. **(iii)** Biochemical parameters and 3D structure of NS1 peptide. **(iv)** Coomassie-stained SDS-PAGE showing purified NS1 at ∼13 kDa and western blot analysis with anti-His. **b) (i)** ELISA**-**based interaction between coated NS1 overlaid with varying concentrations of 14-3- 3□. **c)** Dose-response curve resulting from MST between labeled 14-3-3□ and varying concentrations of NS1. **d)** IFA showing NS1 binding to erythrocyte surface. **e) (i) and (ii)** B3H showing blue and white colonies of the depicted interacting pairs, in the presence and absence of L-arabinose. **f) (i)** Relative MU of the triple co-transformants plotted against varying concentrations of L-arabinose; red line: control and black line: test. **(ii)** Western blot analysis to evaluate the expression of NS1 with increasing L-arabinose concentrations, in R1 cells of triple co-transformants. **g)** Inhibitory ELISA to confirm NS1 mediated inhibition of 14-3-3□/RhopH3 interaction.

### Disruption of 14-3-3**□**/*Pf*RhopH3C interaction results in merozoite invasion inhibition

To investigate the effect of occluding the 14-3-3□/*Pf*RhopH3C interaction upon RBC invasion by the merozoites, three approaches were used: immunological, biochemical, and chemical. In the immunological approach, late schizonts at 1% parasitemia were allowed to invade fresh erythrocytes in the presence of anti-14-3-3□ monoclonal antibody at different dilutions. The newly formed rings were scored and compared with untreated controls to calculate percent invasion inhibition. A dose-dependent reduction in merozoite invasion was observed, with a maximum of reduction ∼85% at a dilution of 1:500 (Fig. 6a). Similarly, in the biochemical approach, schizonts were incubated with uninfected erythrocytes in the presence of 15, 3.75 and 0.78 µM of NS1, and an invasion inhibition of ∼80% was observed in the presence of 15 µM of NS1 (Fig. 6b). In the chemical approach, BV02, a non-peptidyl pyrazolyl carbamoyl inhibitor of 14-3-3, was used. A comparable concentration-dependent reduction in erythrocyte invasion was observed, with a maximum reduction of 80% at 10µM concentration (Fig. 6c). *In silico* interaction analysis validated the binding of RhopH3C, BV02, and NS1 to 14-3-3□, as indicated by negative z-scores (Fig. 6d, e and f). Altogether, this confirms the significance of host 14-3-3□ during the erythrocyte invasion of merozoites.

**Fig. 6:**
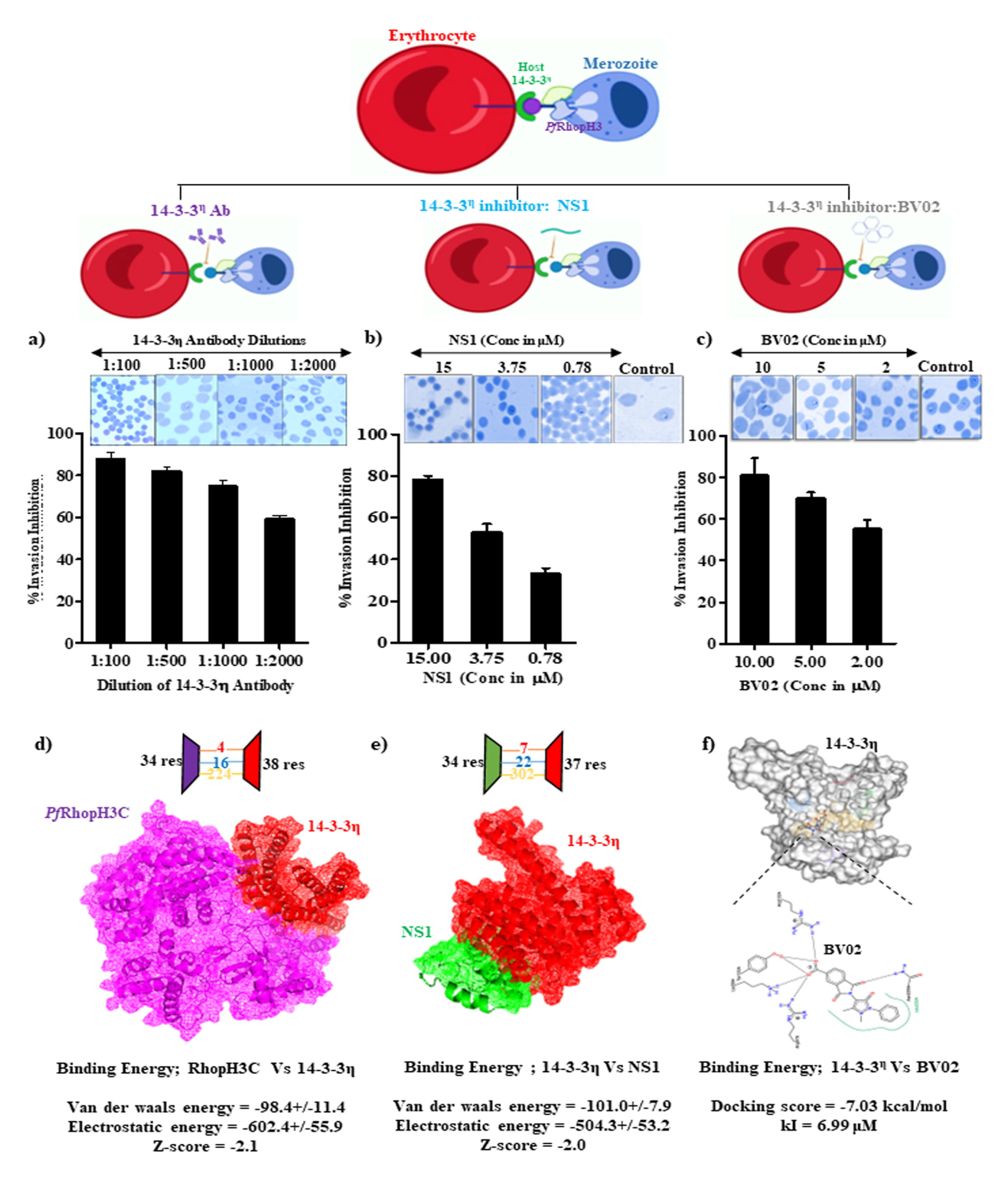
*Pf*3D7 invasion inhibition using immuno-pharmacological approaches. Graph depicting invasion inhibition (%) in the presence of various dilutions of anti-14-3-3□ **(a)**, NS1 **(b)**, and BV02 **(c)**. **d)** Docked complex of 14-3-3□ (red) and *Pf*RhopH3C (magenta). **e)** Docked complex of NS1 (green) and 14-3-3□ (red). Three 14-3-3□ residues were found to be shared in both interactions: LYS-162, ASP-114, and ALA-149. **f)** Blind docking of BV02 and 14-3-3□ showing the participation of ARG132, TYR133, and ASN229 residues from 14-3-3's TBD region.

## DISCUSSION

Secretory proteins are crucial players in host-parasite interactions, rendering them attractive therapeutic targets^32^. Global phospho-proteomic datasets of peptides enriched from the schizont stage of *P. falciparum* are already reported in the literature^33–37^. In this study, we for the first time characterized the protein repertoire secreted by merozoites when exposed to low K^+^-ions, mimicking the conditions encountered by the merozoites in the human body, during erythrocyte invasion. After excluding the major contaminant proteins, a total of 66 proteins were identified, and 50% of them harbored an SP targeting them for secretion. Proteins known to be secreted by the merozoite^38, 39^, such as EBA175, RhopH3, and RAP1 were identified in our secretory fraction, thus validating our findings. GO analysis showed the functional significance of the identified proteins in processes such as protein binding, entry into the host, and symbiotic interaction, and they were found to be associated with the apical region of the parasite which is indispensable for erythrocyte invasion. Furthermore, invasion-related proteins formed a distinct cluster in the secretory fraction that included RhopH3, which was further investigated. After several rounds of B2H with a human cDNA library, we identified 14-3-3□ as a novel interaction partner of *Pf*RhopH3C, which was further validated with multiple biochemical and biophysical approaches.

The 14-3-3 scaffold proteins are known to bind phosphor-ser/thr residues, resulting in structural changes in the associated proteins^38, 40^. Seven isoforms of human 14-3-3 proteins have been identified and disruption in the expression levels of these isoforms has been linked to multiple human disorders including neurodegeneration, parasitic infections, and cancer^41–44^. Recently, the significance of 14-3-3 proteins in SARS-CoV-2 nucleocapsid protein recognition mechanism^45^, and the study of variations in its binding site^46^, provided new insight into their role in host-viral infections^47, 48^. Thus, targeting the host 14-3-3 interactome appears to be an attractive approach for investigating various anti-host-viral/host-parasite strategies. In this study, we for the first time reported the significance of one of the isoforms of human 14- 3-3 protein, eta in malaria pathogenesis.

Identification of 14-3-3 target proteins and their inhibitors/binders that alter disease etiology makes them excellent therapeutic targets. Thus, inhibiting interactions mediated by 14-3-3s could lead to a plethora of therapeutic possibilities. Discovering a novel peptide binder against 14-3-3s to block their interaction with other target proteins without compromising the immune system is a major challenge in this regard. Here, we identified an NS1 peptide that binds to 14-3-3□ and inhibits its interaction with *Pf*RhopH3C, thus depicting a direct involvement of 14-3-3□ and RhopH3C during erythrocyte invasion of merozoites.

All of these findings demonstrated that the interaction between 14-3-3□ and *Pf*RhopH3C is critical for erythrocyte invasion of merozoites. Since the release of rhoptry proteins is important during the invasion, this study establishes a new molecular framework for the development of therapeutics targeting one of the essential rhoptry proteins.

**Declarations of interest:** None.

## Acknowledgements

We are thankful to Central Instrumentation Facility (CIF) of Special Centre for Molecular Medicine, Jawaharlal Nehru University, New Delhi for access to instruments and research facilities. Financial support from Science and Engineering Research Board (SERB) (EMR/2016/005644) to S. Singh is acknowledged. S. Singh is a recipient of the IYBA Award from DBT. This work is also supported by the Science and Engineering Research Board under the Department of Science and Technology (CRG/2019/002231, IPA/2020/000007) sanctioned to AR and S. Singh. The funders had no role in study design, data collection, and analysis, decision to publish, or preparation of the manuscript.

## Author contribution

The study was conceptualized and designed by S. Singh, A.R. and P.M. Acquisition of data was done by N.S., A.M., G.K., S.K., A.K.K., S.G., A.C., Z.A., M.K.M., P.S., J.S., M.M., P.J., R.J., D.A.B.R., T.S.K.P., and M.M. Data analysis and interpretation was done by N.S., A.M., G.K., S.K., S. Singh and A.R. Manuscript was written, drafted and revised by N.S., A.M., R.J., S. Singh and A.R. All authors approved final version of the manuscript.

